# Extracting Phylogenetic Information of Human Mitochondrial DNA by Linear Autoencoder

**DOI:** 10.1101/2021.06.22.449384

**Authors:** Takuto Shimazaki, Kazuki Nishimoto, Hirohiko Niioka, Jun Miyake

**Affiliations:** Department of Material and Life Science, Graduate School of Engineering, Osaka University, Suita, Osaka, Japan; School of Engineering Science, Osaka University, Suita, Osaka, Japan; Graduate School of Information Science and Technology, Osaka University, Suita, Osaka, Japan; Institute for Datability Science, Osaka University, Suita, Osaka, Japan; Hitz Research Alliance Laboratory, Graduate School of Engineering, Osaka University, Suita, Osaka, Japan; Osaka University, Global Center for Medical Engineering and Informatics, Suita, Osaka, Japan

**Keywords:** Principal component analysis, Linear autoencoder, Gradient descent, Human, Mitochondrial DNA, Phylogenetic tree, Genome

## Abstract

We used a linear autoencoder (LAE) and its learning dynamics to analyze the high-order structure of human mitochondrial DNA (mtDNA). A total of 360 complete human mtDNA sequences were collected from the MITOMAP database and transformed into 1024-dimensional vectors of pentanucleotide frequencies. We compressed those into a three-dimensional (3D) coordinates by an LAE at each step of training by gradient descent with respect to the quadratic error function. Along the time axis of training epochs, the compressed 3D coordinates were gradually clustered and separated in accordance with the order of the genetic distance in the phylogenetic tree of human mtDNA haplogroups. This suggests that there is an association between the learning dynamics of LAE and the high-dimensional structure of human mtDNA sequences, similar to that of phylogenetic analysis and evolutionary pathways: the five clusters eventually contained only a single haplogroup of L0, M, N, R, and U, while the L3 cluster contained a small number of M members and The packing was comparable to that realized in learning dynamics similar to genetic classification and evolutionary pathways by LAE in principal component analysis (PCA), but somewhat denser than PCA.

## Introduction

In population genetics, the application of principal component analysis (PCA) has widely been explored since it was first introduced to the analysis of genetic diversity within the human genome in the 1970s (Menozzi *et al*. 1978). PCA effectively summarizes high-dimensional genomic data by projecting them onto a low-dimensional space, enabling us to analyze major genetic patterns visually. A wide range of genetic structures has been uncovered by PCA, such as the correspondence of genetic relatedness with the geographic distances among Europeans (Novembre *et al*. 2008) and the influence of genealogical history on PCA projections (McVean 2009; Patterson *et al*. 2006). However, identifying genetic patterns from PCA projections remains a major issue owing to the non-parametric feature of PCA. Also, non-genetic trends, such as statistical artifacts that do not necessarily reflect the high-order structure, can confound the interpretation of PCA projection (Novembre & Stephens 2008). To improve the interpretability of PCA, a method that effectively extracts the high-order structure of genomic data and is still communicable with PCA projections is necessary.

In the field of machine learning, it is known that linear autoencoders (LAEs) and PCA are closely related algorithms; an LAE trained to minimize the quadratic error function also learns the subspace spanned by the top principal eigenvectors of the covariance matrix of given training data (i.e., the first eigenvectors of the covariance matrix in descending order by eigenvalues) (Bourlard & Kamp 1988; Baldi & Hornik 1989). Autoencoders are a dimensionality reduction algorithm used to extract features of high-dimensional data. An autoencoder compresses and recovers high-dimensional input through its encoding and decoding processes and update its parameters so that the reconstructed data match the original input. The optimal parameters are usually found by running a gradient method, such as gradient descent, on an error function, which measures how well the reconstructed data represent the original input. As the optimization progresses, the compressed low-dimensional data are better represented and can be regarded as a summary of the high-dimensional input. Unlike PCA, most autoencoders are nonlinear in that they are equipped with nonlinear functions, called activation functions, in the encoding and decoding parts, and the nonlinearity is said to capture the nonlinear features of high-dimensional data (Hinton and Salakhutdinov 2006). LAEs refer to the autoencoders that have identity functions (*f*(*x*) = *x*) for the activation functions. When an LAE is trained to minimize the quadratic error function, the linearization of the activation functions dramatically simplifies the landscape of the quadratic error function as follows (Bourlard & Kamp 1988; Baldi & Hornik 1989; Kunin *et al*. 2019).

i. The quadratic error function has a unique local and global minimum up to invertible linear transformations, and all the other critical points are saddle points.
ii. The unique minimum of the quadratic error function corresponds to the projection onto the subspace spanned by the top principal eigenvectors, and the saddle points correspond to the projections onto the subspaces spanned by principal eigenvectors of the other combinations.

The two mathematical properties above with a further mathematical consideration imply that, when an LAE is trained by gradient descent, the local minimum-free landscape of the quadratic error function guarantees the convergence to its unique minimum (Kunin *et al*. 2019). This informs that the learning dynamics of LAEs induced by gradient descent generate a series of continuous projections at each training epoch that eventually reaches the projection equivalent to PCA. In this study, we demonstrate that the continuous projections characterized by the learning dynamics of LAE can translate back to genetic meaning and potentially provide further insight into the population structure inference by PCA.

## Materials and Methods

We briefly describe the vectorization method for the human mitochondrial DNA (mtDNA) sequential data, the algorithm and the optimization of LAEs used in this study. Further details for the vectorization method and the LAE algorithm can be found in Müller & Koonin (2003) and Baldi & Hornik (1989), respectively (Müller & Koonin 2003; Baldi & Hornik 1989).

A total of 360 complete human mtDNA sequences with 16560 nucleotides (nt), consisting of six haplogroups L0, L3, M, N, R, and U with 60 sequences for each, were obtained from the MITOMAP database (Kaneshita *et al*. 2015). We transformed the sequences into fixed-size vectors by counting the occurrences of words called n-grams (Fig. 1). N-gram-based vectorizations are frequently used in the field of computational linguistics and also shown to be effective in genomic data analysis, such as the classification of introns, exons, and intergenic regions (Müller & Koonin 2003) and the classification of eukaryotic genomes (Abe *et al*. 2002; Abe *et al*. 2003). A word of size *l* is defined as a string of *l* consecutive characters, consisting of A, T, C, or G. We count the occurrences of words, *K_i_* (*i* indicates the index for a particular word), in a sequence by moving a window of size *l* character by character along the sequence allowing overlaps. This process produces a vector whose components are the occurrences of words. The size of the vector becomes 4^*l*^ as there are 4^*l*^ combinations of words in the case of genomic sequences. In this study, the value of *l* was heuristically determined as 5 (Miyake *et al*. 2018), resulting in the vector size of 1024. We applied this method to all of the 360 human mitochondrial sequences and created a 360 ×1024 matrix by combining the vectors by rows. This matrix was regarded as the summary of the pentanucleotide frequencies of the human mtDNA sequences and used as an input to an LAE.

**Fig. 1.**
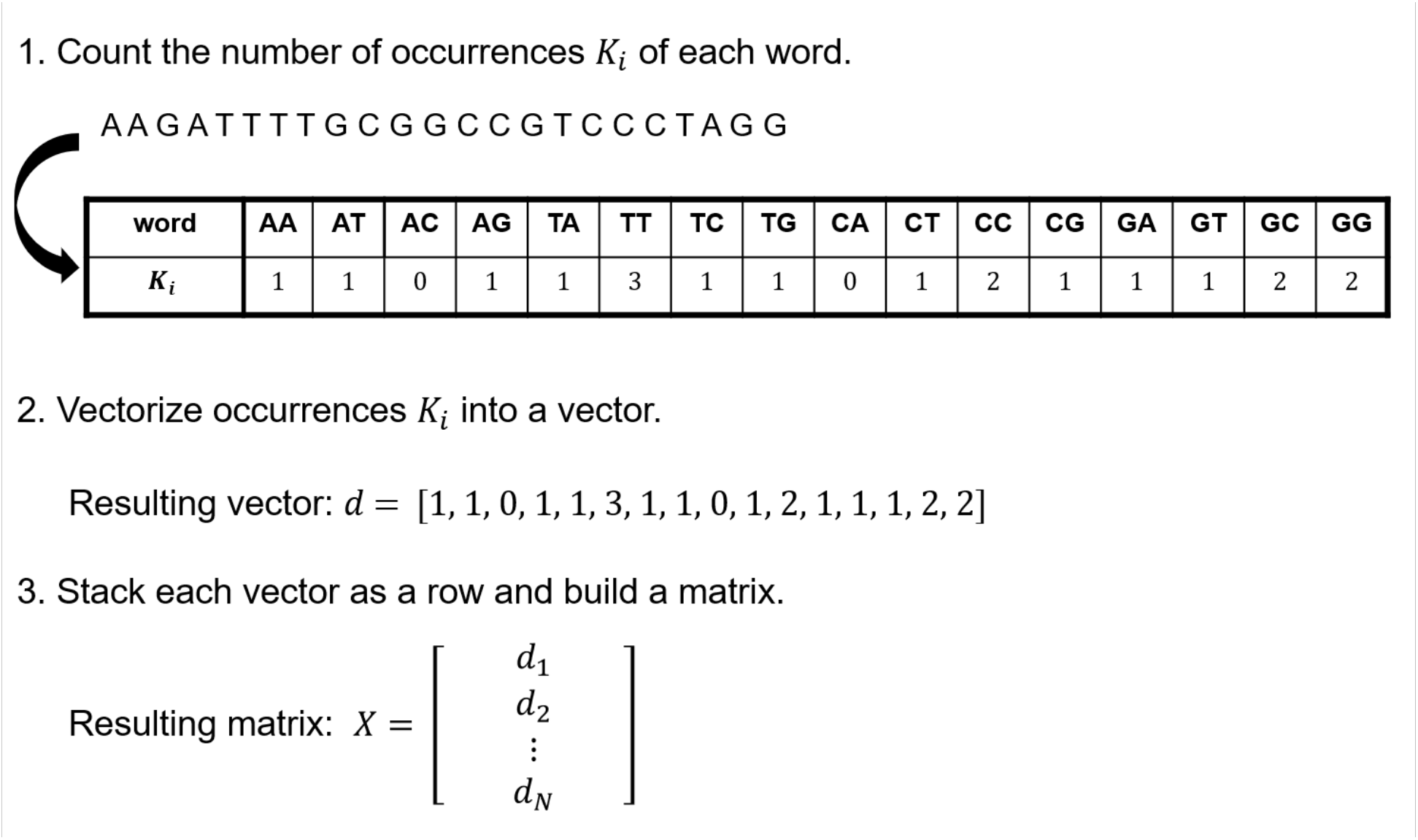
Transformation of genomic sequences into a matrix of word frequencies. This figure shows three steps to transform a genetic sequence of arbitrary length into a fixed-size vector. In the example above, the length of a word *l* is set to 2, resulting in 4^2^ = 16 possible words and vectors size of 16.

An LAE consists of an encoder and a decoder; an encoder compresses the high-dimensional input data, and a decoder reconstructs the original input from the compressed data. Denoting the number of the sequences by ***N*** and the dimensions of the original and compressed data by ***m*_1_** and ***m*_2_**, respectively, the compressed data, 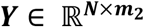, is expressed by

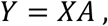

where 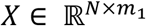 is the input matrix, and 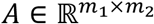 is the weight matrix constituting the encoder. The reconstructed data, 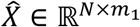, is expressed by

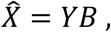

where 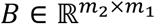 is the weight matrix constituting the decoder. In this study, we used the quadratic error function to make the autoencoder learn the principal subspace. The quadratic error function is defined by

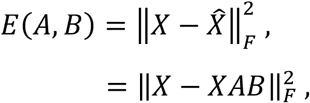

where ∥ • ∥_*F*_ denotes the Frobenius norm. In the training process, the weight matrices *A* and *B* are updated by gradient descent to minimize the quadratic error function. Let *A*(*t*) and *B*(*t*) be the weight matrices *A* and *B* after *t* times of updates by gradient descent and *a_ij_*(*t*) and *b_ij_*(*t*) be the entry in the *i*th row and *j*th column of *A*(*t*) and *B*(*t*). By gradient descent, *a_ij_*(*t*) and *b_ij_*(*t*) are updated to *a_ij_*(*t* + 1) and *b_ij_*(*t* + 1) by

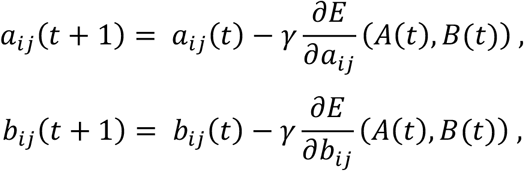

where *γ* (> 0) is the learning coefficient, which adjusts the step size of updates. In the formulation above, we omitted bias parameters assuming that the input data are mean-centered. This formulation with the assumption is equivalent to the one with bias parameters (Baldi and Hornik 1989; Kunin *et al*. 2019). Also, we restricted the formulation to single-layer LAEs because an LAE with more than one hidden layer reduces to the case of a single-layer LAE owing to its linearity (Baldi & Hornik 1989; Kunin *et al*. 2019).

## Results

The 360 human mtDNA sequences of the haplogroups L0, L3, M, N, R, and U were vectorized and represented as a 360 × 1024 matrix. After mean-centering the matrix, an LAE was trained for a sufficient number of epochs by gradient descent to minimize the quadratic error function, and the vectorized human mtDNA sequences were projected to the subspace learned at the 300th, 1200th, 1700th, 3000th, 7000th, and 10,000th epochs (Fig. 2A–F). Owing to the local minimum-free landscape, the value of the quadratic error function monotonically decreased with training epochs until it converged to a certain value approximately at the 7000th epoch (Fig. 2G).

**Fig. 2.**
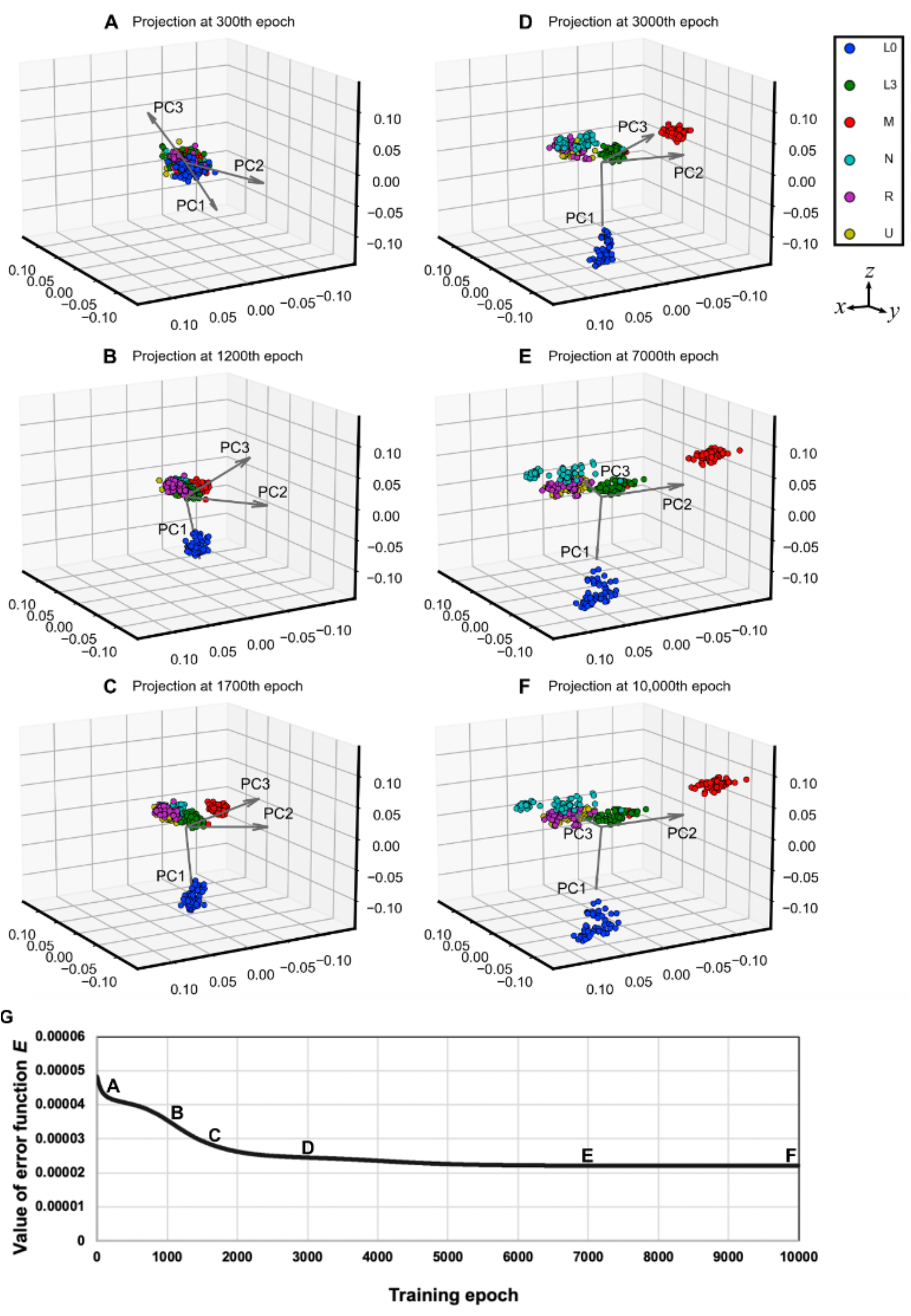
Phylogenetic separations projected by an LAE at six training epochs and the value of the quadratic error function along the training epochs. (**A**) A total of 360 human mtDNA sequences were projected onto the three-dimensional subspace learned by an LAE at the 300th epoch. (**B**) the 1200th epoch. (**C**) the 1700th epoch. (**D**) the 3000th epoch. (**E**) the 7000th epoch. (**F**) the 10,000th epoch. The dots in each projection represent the projected sequences, and the dot colors represent different human mtDNA haplogroups (L0, blue; L3, green; M, red; N, cyan; R, violet; U, yellow). (**G**) The values of the quadratic error function along the training epochs. Suffixes were placed to associate the projections with the corresponding points on the error curve. The three gray arrows in (**A**)–(**F**) indicate the principal directions in the 1024-dimensional space projected onto the threedimensional space learned by the LAE at the displayed training epochs. PC1, PC2, and PC3 arrows (equal in length) mean the first, second, and third principal directions, respectively, projected onto the three-dimensional space learned by the LAE at respective training epochs. Full video of the cluster transition from the 1st epoch to the 10,000th epoch is available at *https://www.miyakelab.com/research*.

At the 300th epoch, when the values of weight matrices only slightly differ from the randomly initialized values, the compressed 3D coordinates (represented by dots) of human mtDNA sequences were in a mixed state around the center (Fig. 2A). From the 300th epoch to the 1200th epoch, the dots corresponding to the sequences of the L0 clade were smoothly separated from the original group at the center and newly formed a new distinct cluster consisting of the L0 clade exclusively (Fig. 2B). Soon after the separation of the L0 clade, the dots corresponding to the M clade followed a gradual separation along the perpendicular direction to the L0 dissociation direction (Fig. 2C). Next, toward the opposite direction of the M-clade separation, the dots for the N, R, and U clades were separated together to form a mixed cluster of the N, R, and U clades (Fig. 2D). These M-clade and subsequent N-, R-, and U-clades separations happened almost in parallel from the 1200th epoch to the 3000th epoch. From the 3000th epoch, the dots corresponding to the N, R, and U clades showed a separation into two distinct clusters of the N clade and the R and U clades, and the cluster of the R and U clades was further subdivided into two U-clade and one R-clade clusters, with the former two positioned beside the latter (Fig. 2E). No notable changes were observed in the cluster distribution after the 7000th epoch (Fig. 2F). Remaining L3 cluster included a few members of the M clade.

To analyze the learning behavior of the LAE, we projected the top three principal directions onto the 3D space learned by the LAE at each training epoch and expressed them as arrows (Fig. 2). During the training of the LAE, the transition of the arrow directions was characterized by three phases of distinct gradients of the error function: A (from the 0th epoch to the 1200th epoch), B (from the 1200th epoch to the 3000th epoch), and C (from the 3000th epoch to the 7000th epoch). As a whole, the LAE learned the principal directions in descending order. In phase A, the PC1 arrow adjusted its direction toward the L1-clade separation before the separation happened, with the PC2 arrow slowly changing its direction toward the subsequent M-clade separation and the PC3 arrow showing a somewhat random behavior. Phase B was triggered near the end of the L1-clade separation, and the PC2 arrow was fixed to the direction of the M-clade separation, with the PC1 arrow keeping its direction and the PC3 arrow gradually changing its direction of the subsequent N-, R-, and U-clade separations. During phase C, all the PC arrows held their directions, with the PC3 arrow pointing to the N-, R-, and U-clade separations. One feature common to all the phases was that each phase occurred concurrently with the major separations of the projected human mtDNA sequences.

As mentioned in the introduction, the PCA projection and the projection by a converged LAE trained to minimize the quadratic error function is the same up to deformation by invertible linear transformations. Comparing the LAE and PCA projections in the case of the 360 human mtDNA sequences led to observations corroborating the prediction by theory; both projections showed the same pattern of cluster separations and the relative cluster positions, with dissimilarities predominantly arising from slightly more dense packing in respective clusters (LEAs), rotations and scalar transformations (Fig. 3).

**Fig. 3.**
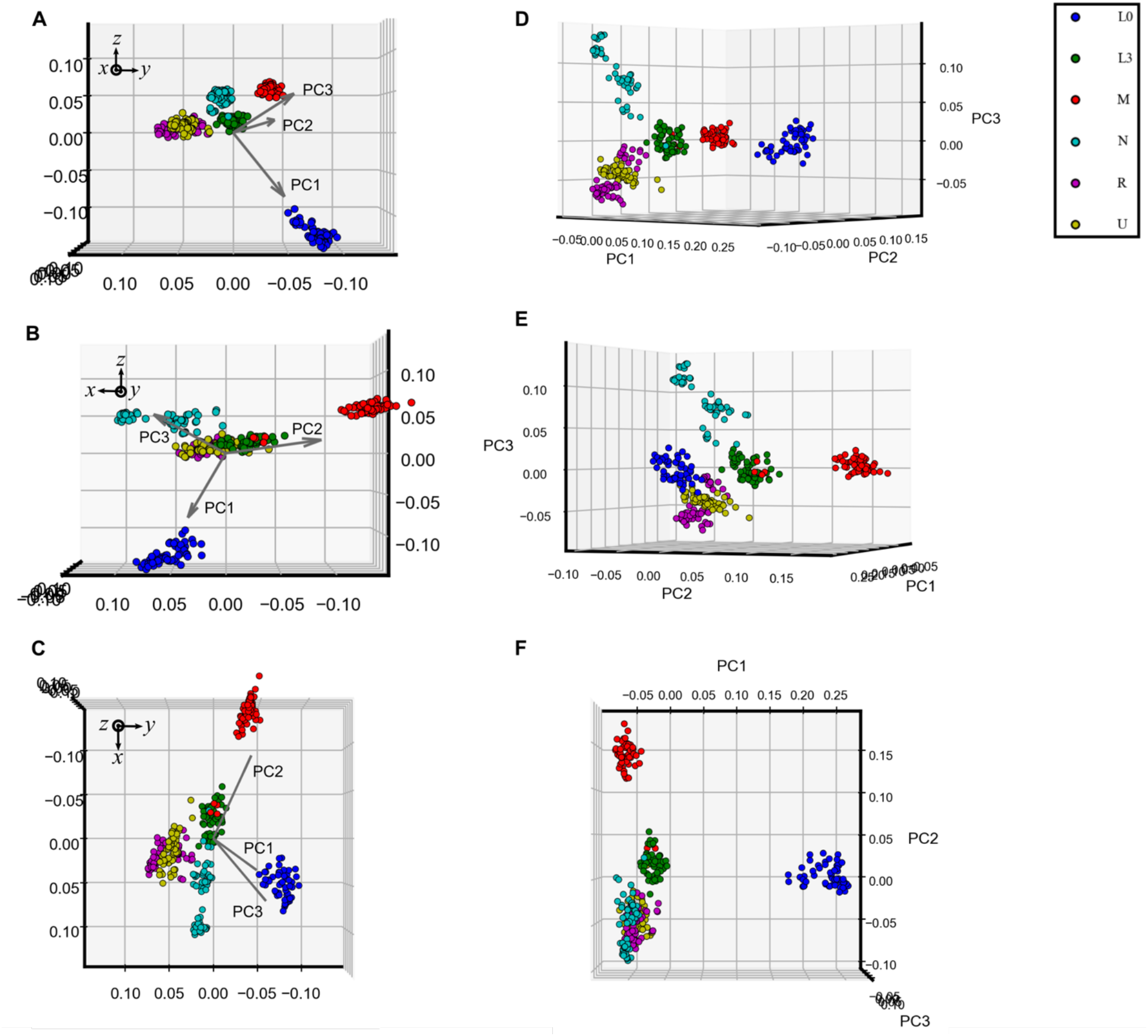
Comparison of LAE and PCA projections. (**A**) The 360 human mtDNA sequences were projected by a fully trained LAE and were seen from the x-axis. (**B**) from the y-axis. (**C**) from the z-axis. (**D**) The 360 human mtDNA sequences were projected by PCA and were seen from an angle similar to (**A**). (**E**) seen from an angle similar to (**B**). (**F**) from an angle similar to (**C**). The x, y, and z directions match the ones in Fig. 2. The dots are colored as described in Fig. 2.

## Discussion

Considering that mutations along maternal lineages determine the variation of human mtDNA sequences (Torroni *et al*. 2006), a set of mtDNA sequences may form a tree-like structure in the high-dimensional space, whose branches representing each mutation radiate to various dimensions. To support this, the observed cluster separations along the training epochs showed a strong correlation with the branching pattern of phylogenetic tree of human mtDNA haplogroups (Fig. 4) (Shinoda 2019). This implies that the LAE learned the tree-like structure in a manner that follows the phylogenetic tree from its origin. In addition to genetic classification, this study also exemplifies the performance of LEAs for showing evolutionary pathway-like learning dynamics of LAEs, being advantageous to PCA.

**Fig. 4.**
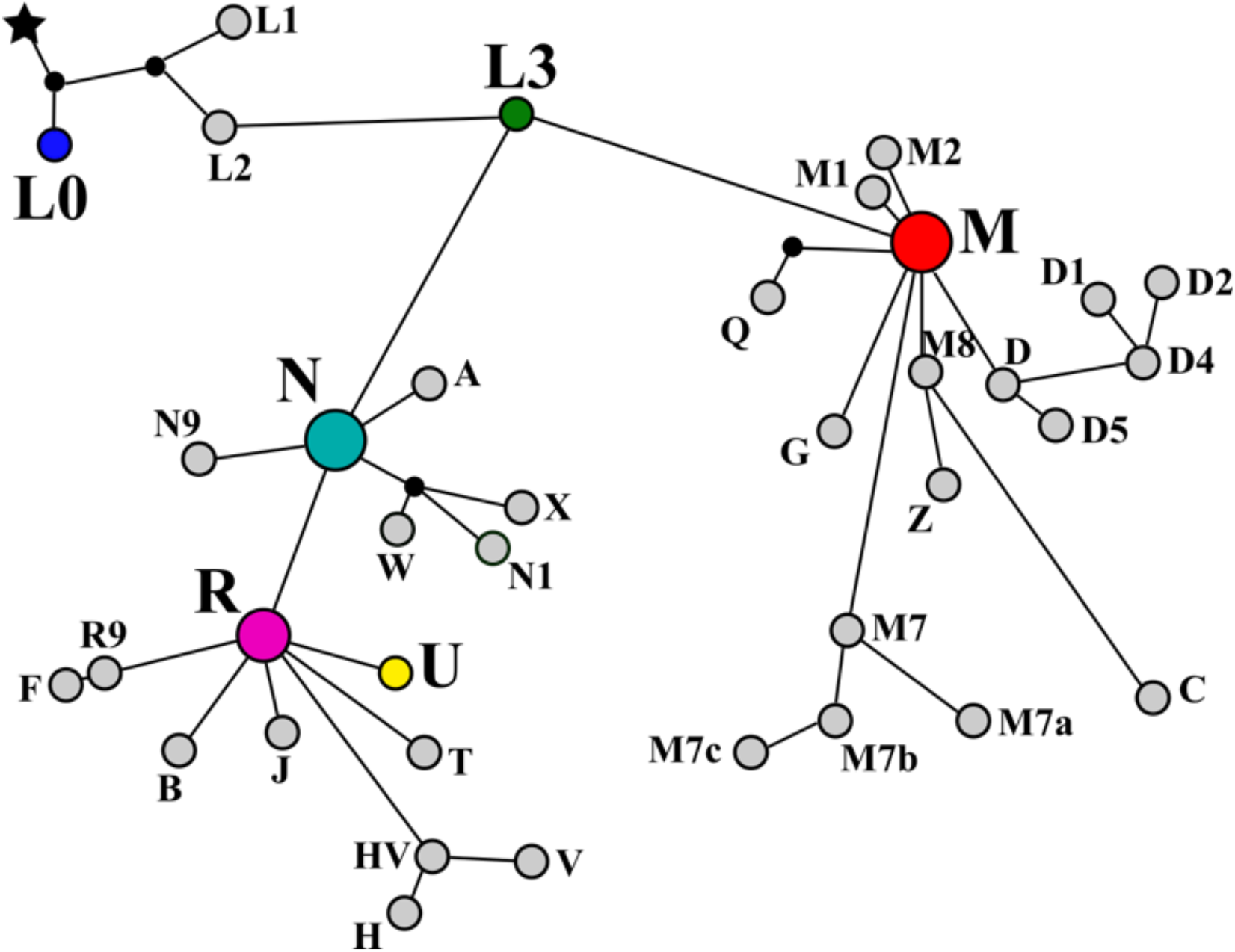
Phylogenetic tree of human mtDNA haplogroups. This figure indicates the phylogenetic relationship human mtDNA haplogroups. The nodes of haplogroups L0, L3, M, N, R, and U are colored as described in Fig. 2. This figure was adapted from the paper by Shinoda (Shinoda 2019).

The phylogenetic separations of human mtDNA sequences can be explained as follows. The projections of the top three principal directions imply that the LAE learns the top principal directions in descending order, one by one (Fig. 2). Since the principal directions are made up of a linear combination of pentanucleotide frequencies, which are characterized by mutations, the learning dynamics acted on the LAE to distinguish the sequences with a specific set of mutations and the ones without it, from the mutations that accounted for the largest genetic variation. In other words, the LAE gradually changed the direction of the axes of the low-dimensional subspace towards the steepest direction on the landscape of the quadratic error function creating silhouettes on the low-dimensional subspace, which appeared as a single cluster at first and separated as they follow the phylogenetic relationships along the time of evolution.

LAEs do not necessarily capture the entire phylogenetic relationships as evident from the fact that LAEs learned with a quadratic error function converges to a projection equivalent to PCA; their projections only reflect the genetic variation in the learned low-dimensional subspace. To show an example, we trained an LAE that learns a two-dimensional subspace for the same data and projected the sequences onto the learned subspace after convergence (Fig. 5). As a result, the expansion along the N-, R-, and U-clades observed in the case of 3D subspace collapsed into a single cluster, suggesting that the LAE failed to capture the genetic variation among the N, R, and U clades. To investigate further details of genetic variation, we need to consider other methods, such as running PCA locally with a smaller sample size and nonlinear models.

**Fig. 5.**
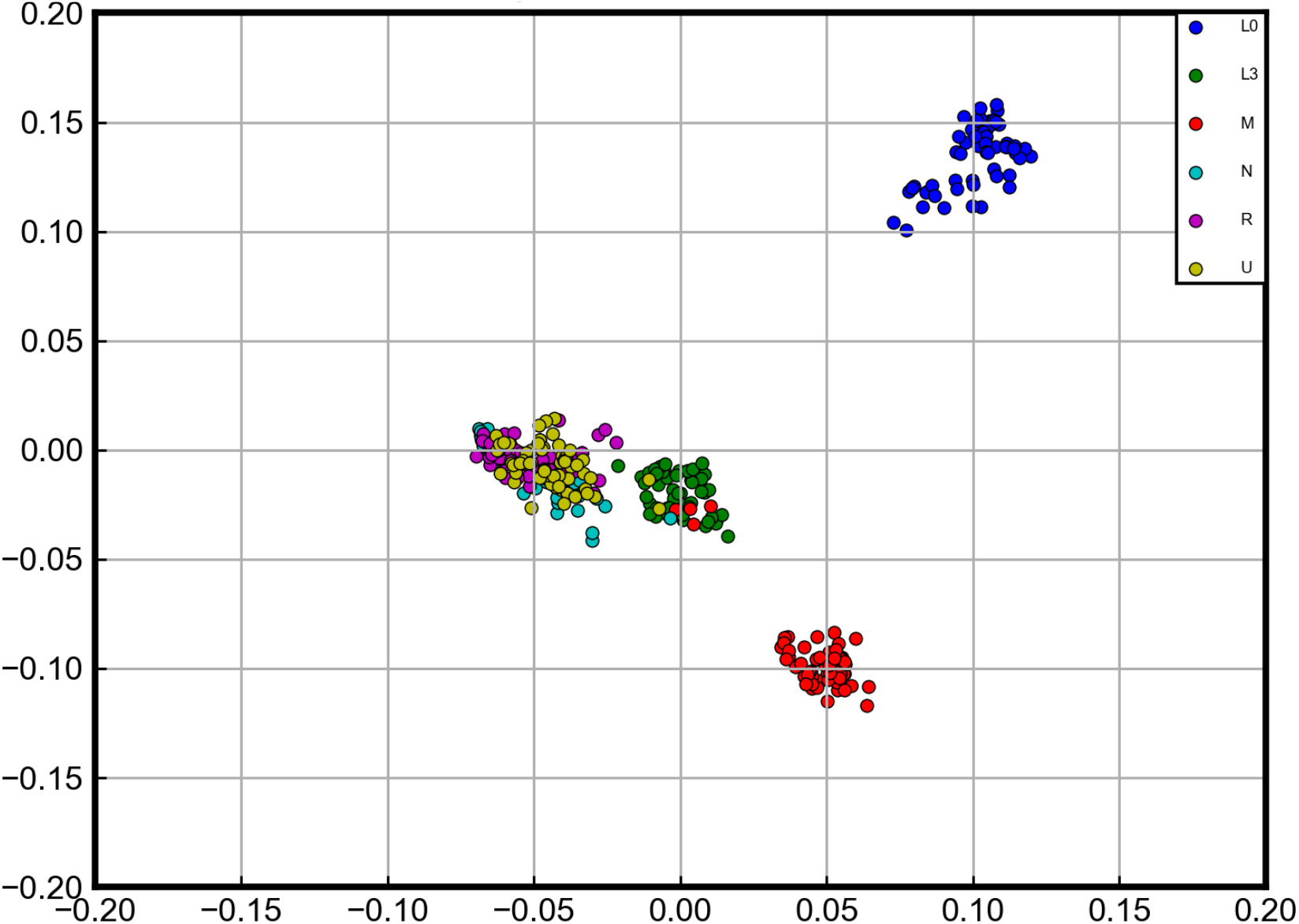
Projection onto the two-dimensional subspace by a trained LAE. A total of 360 human mtDNA sequences were projected onto the two-dimensional subspace learned by an a fully trained LAE. The dots are colored as described in Fig. 2. Note that, unlike the case of the projection onto three-dimensional subspace, the clusters of N, R, and U clades collapsed into a single cluster.

Several studies have reported the correlation between phylogenetic relationships in genomic data and projections by dimensionality reduction algorithms, such as PCA (Alexe *et al*. 2008; Miyake *et al*. 2021). Those studies suggest that LAEs can learn phylogenetic relationships from genomic data that can be clustered based on mutations. The extent to which the present results generalize to other genomic data is a meaningful issue.

This work exemplifies the genetic variation of genomic data is not only explained by PCA, which corresponds to the unique minimum on the landscape of the quadratic error function, but the entire landscape can also be subject to analysis and provide a further insight into the population structure inference by PCA. To the best of our knowledge, this study is the first attempt to extract biological meaning from genomic data by utilizing the dynamics on the landscape

## Author contributions

TS originated the project; TS and KN wrote the manuscript; TS developed computer system and software; HN supervised the study of machine learning; JM supervised and wrote the work.

## Acknowledgements

We should like to express our thanks to Takaaki Sato and Hayato Nakamura of our laboratory for reviewing the work. The authors express special thanks to Prof. Yoshihisa Nakazawa of Hits Research Alliance Laboratory of Osaka University for his encouragements and supports. This work was supported partially by Global Center for Medical Engineering and Informatics of Osaka University, Hitz Research Alliance Laboratory of Osaka University and Nihon Unisys, Ltd.

## References

Abe T, Kanaya S, Kinouchi M, Ichiba Y, Kozuki T, Ikemura T. A novel bioinformatic strategy for unveiling hidden genome signature of eukaryotes: self-organizing map of oligonucleotide frequency. Genome Inform. 13:12–20 (2002). doi: 10.11234/gi1990.13.12.

Abe T, Kanaya S, Kinouchi M, Ichiba Y, Kozuki T, Ikemura T. Informatics for unveiling hidden genome signatures. Genome Res. 13:693–702 (2003). doi: 10.1101/gr.634603.

Alexe G, Satya RV, Seiler M, Platt D, Bhanot T, Hui S, Tanaka M, Levine AJ, Bhanot G. PCA and clustering reveal alternate mtDNA phylogeny of N and M clades. J Mol Evol. 67:465–487 (2008). doi: 10.1007/s00239-008-9148-7.

Baldi P, Hornik K. Neural networks and principal component analysis: learning from examples without local minima. Neural Netw. 2:53–58 (1989). doi: 10.1016/0893-6080(89)90014-2.

Baldi P, Hornik K. Learning in linear neural networks: a survey. IEEE Trans Neural Netw. 6:837–858 (1995). doi: 10.1109/72.392248.

Bourlard H, Kamp Y. Auto-association by multilayer perceptrons and singular value decomposition. Biol Cybern 59:291–294 (1988). doi: 10.1007/BF00332918.

Cavalli-Sforza LL, Feldman WM. The application of molecular genetic approaches to the study of human evolution. Nat Genet. 33:266–275 (2003). doi: 10.1038/ng1113.

Goodfellow I, Bengio Y, Courville A. Deep learning. MIT Press (2016).

Hinton EG, Salakhutdinov RR. Reducing the dimensionality of data with neural networks. Science 313:504–507 (2006). doi: 10.1126/science.1127647.

Ingman M, Kaessmann H, Pääbo S, Gyllensten U. Mitochondrial genome variation and the origin of modern humans. Nature 408:708–713 (2000). doi: 10.1038/35047064.

Kaneshita Yuhei, Sugiyama Kanako, Asatani Satoshi, Niioka Hiirohiko, Hirano Takashi, Miyake Jun, Classification of Mitochondrial DNA with using deep learning, Proceedings of 16th SICE System Integration Division Annual Conference: 1267–1270 (2015).

Kunin D, Bloom J, Goeva A, Seed C. Loss landscapes of regularized linear autoencoders. Proceedings of the 36th International Conference on Machine Learning, PMLR 97:3560–3569 (2019).

McVean G. A genealogical interpretation of principal components analysis. PLoS Genet. 5:e1000686 (2009). doi: 10.1371/journal.pgen.1000686.

Menozzi P, Piazza A, Cavalli-Sforza L. Synthetic maps of human gene frequencies in Europeans. Science 201:786–792 (1978). doi: 10.1126/science.356262.

Menozzi P, Cavalli-Sforza LL, Piazza A. Demic expansions and human evolution. Science 259:639–646 (1993). doi: 10.1126/science.8430313.

Miyake J, Kaneshita Y, Asatani S, Tagawa S, Niioka H, Hirano T. Graphical classification of DNA sequences of HLA alleles by deep learning. Hum Cell. 31:102–105 (2018). doi: 10.1007/s13577-017-0194-6.

Miyake J, Sato T, Baba S, Nakamura H, Niioka H, Nakazawa Y. Cluster analysis of SARS-CoV-2 gene using deep learning autoencoder: gene profiling for mutations and transitions. BioRxiv (2021). doi: 10.1101/2021.03.16.435601.

Muller H-M, Koonin SE. Vector space classification of DNA sequences. J Theor Biol. 223:161–169 (2003). doi: 10.1016/S0022-5193(03)00082-1.

Nielsen R, Akey MJ, Jakobsson M, Pritchard KJ, Tishkoff S, Willerslev E. Tracing the peopling of the world through genomics. Nature 541:302–310 (2017). doi: 10.1038/nature21347.

Novembre J, Johnson T, Bryc K, Kutalik Z, Boyko RA, Auton A, Indap A, King SK, Bergmann S, Nelson RM, Stephens M, Bustamante DC. Genes mirror geography within Europe. Nature 456:98–101 (2008). doi: 10.1038/nature07331.

Novembre J, Stephens M. Interpreting principal component analyses of spatial population genetic variation. Nat Genet. 40:646–649 (2008). doi: 10.1038/ng.139.

Novembre J, Ramachandran S. Perspectives on human population structure at the cusp of the sequencing era. Annu Rev Genomics Hum Genet. 12:245–274 (2011). doi: 10.1146/annurev-genom-090810-183123.

Oja E. Neural networks, principal components, and subspaces. Int J Neural Syst. 1:61–68 (1989). doi: 10.1142/S0129065789000475.

Patterson N, Price A, David Reich. Population structure and eigenanalysis. PLoS Genet. 2:e190 (2006). doi: 10.1371/journal.pgen.0020190.

Plaut E. From principal subspaces to principal components with linear autoencoders. arXiv: 1804.10253 (2018). https://arxiv.org/pdf/1804.10253

Shinoda K. Analysis of DNA variations reveals the origins of dispersal of modern humans. J Geog. 118:311–319 (2009). doi: 10.5026/jgeography.118.311.

Torroni A, Achilli A, Macaulay V, Richards M, Bandelt H-J. Harvesting the fruit of the human mtDNA tree. Trends in Genet. 22:339–345 (2006). doi: 10.1016/j.tig.2006.04.001.

